## Additional File 2 for "Predicting yield traits of individual field-grown *Brassica napus* plants from rosette-stage leaf gene expression"

### Additional File 2: Figures S1-S11

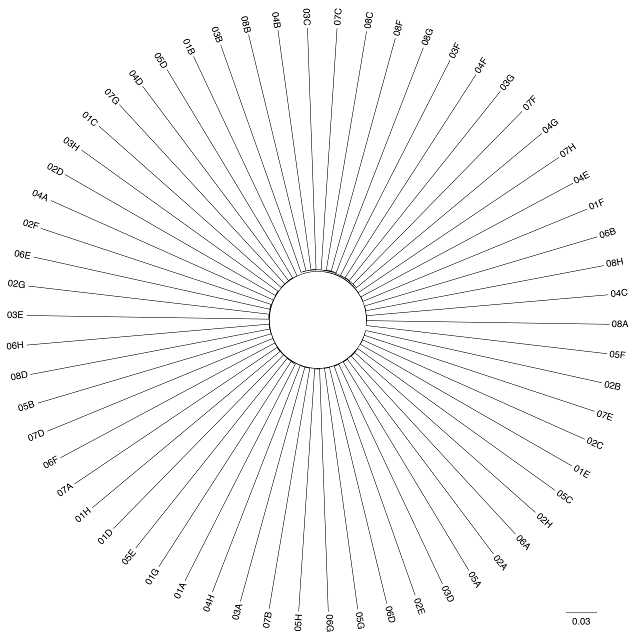


**Fig. S1** SNP analysis of individual plants. Neighbor-joining tree for the individual plants based on biallelic SNPs in the RNA-seq data. Branch lengths are proportional to 1 – IBS (identity by state).

**Fig. S2.** Phenotype field plots. Each plot displays the variation of a phenotype over the field. Shown on top of each plot are the Moran's I value of the phenotype, the expected Moran's I (by resampling) and the associated *p*-value and *q*-value (computed using the Benjamini-Hochberg method over all phenotypes). The table below orders phenotypes by increasing *p*-value, the page and panel numbers of each plot are given in the 'page' and 'panel' columns, respectively.

| phenotype | Moran's I | p-value | q-value | page | panel |
|---|---|---|---|---|---|
| root system width | -0.037026 | 0.00061 | 0.025010 | 6 | D |
| rosette area (42 DAS) | 0.001414 | 0.00931 | 0.190855 | 2 | A |
| plant height (278 DAS) | 0.269224 | 0.01438 | 0.196527 | 8 | D |
| seed weight stem 1/dry weight stem 1 | 0.010629 | 0.02511 | 0.242966 | 6 | F |
| total seed weight/shoot dry weight | 0.022452 | 0.02963 | 0.242966 | 6 | E |
| branch count stem 1/length stem 1 | 0.031889 | 0.04530 | 0.280791 | 4 | F |
| rosette lesions (74 DAS) | 0.225559 | 0.04794 | 0.280791 | 2 | E |
| leaf 8 lesions (76 DAS) | 0.192259 | 0.11757 | 0.462517 | 3 | C |
| siliques per branch | 0.192239 | 0.11762 | 0.462517 | 6 | A |
| taproot length | 0.191263 | 0.12236 | 0.462517 | 6 | C |
| leaf 6 length (74 DAS) | 0.189472 | 0.12409 | 0.462517 | 2 | B |
| siliques per branch stem 1 | 0.179719 | 0.16236 | 0.495170 | 6 | B |
| max shoot growth rate | 0.179163 | 0.17559 | 0.495170 | 8 | A |
| silique count stem 1 | 0.170856 | 0.19550 | 0.495170 | 5 | F |
| leaf 8 chlorophyll content (81 DAS) | 0.169002 | 0.22343 | 0.495170 | 3 | D |
| total branch count | 0.077732 | 0.22679 | 0.495170 | 4 | C |
| dry weight stem 1 | 0.162020 | 0.23601 | 0.495170 | 5 | B |
| dry weight stem 1 (w/o seeds) | 0.159648 | 0.24895 | 0.495170 | 5 | D |
| leaf 8 area (81 DAS) | 0.154198 | 0.27305 | 0.495170 | 3 | E |
| seed weight stem 1 | 0.150570 | 0.29601 | 0.495170 | 7 | F |
| seed count stem 1 | 0.148525 | 0.31013 | 0.495170 | 7 | B |
| leaf 8 fresh weight (81 DAS) | 0.147416 | 0.31125 | 0.495170 | 4 | A |
| end of shoot growth | 0.095732 | 0.31765 | 0.495170 | 8 | B |
| leaf 8 length (81 DAS) | 0.144311 | 0.32831 | 0.495170 | 3 | F |
| seeds per silique | 0.101385 | 0.35827 | 0.495170 | 7 | C |
| branch count stem 1 | 0.099686 | 0.36557 | 0.495170 | 4 | D |
| time of max shoot growth | 0.106377 | 0.38642 | 0.495170 | 8 | C |
| total shoot dry weight (w/o seeds) | 0.132044 | 0.41698 | 0.495170 | 5 | C |
| leaf 8 width (81 DAS) | 0.129682 | 0.41995 | 0.495170 | 4 | B |
| leaf 8 width (76 DAS) | 0.129013 | 0.42559 | 0.495170 | 3 | B |
| stem count | 0.109016 | 0.43006 | 0.495170 | 8 | E |
| total shoot dry weight | 0.128425 | 0.43706 | 0.495170 | 5 | A |
| leaf 6 width (74 DAS) | 0.125089 | 0.45185 | 0.495170 | 2 | C |
| total seed count | 0.112390 | 0.45234 | 0.495170 | 7 | A |
| leaf count (74 DAS) | 0.124303 | 0.45869 | 0.495170 | 2 | D |
| seeds per silique stem 1 | 0.124687 | 0.46503 | 0.495170 | 7 | D |
| leaf 8 length (76 DAS) | 0.122295 | 0.47394 | 0.495170 | 3 | A |
| total seed weight | 0.115616 | 0.47672 | 0.495170 | 7 | E |
| leaf 6 lesions (74 DAS) | 0.115471 | 0.48131 | 0.495170 | 2 | F |
| total silique count | 0.121435 | 0.48501 | 0.495170 | 5 | E |
| branches per stem | 0.119628 | 0.49517 | 0.495170 | 4 | E |

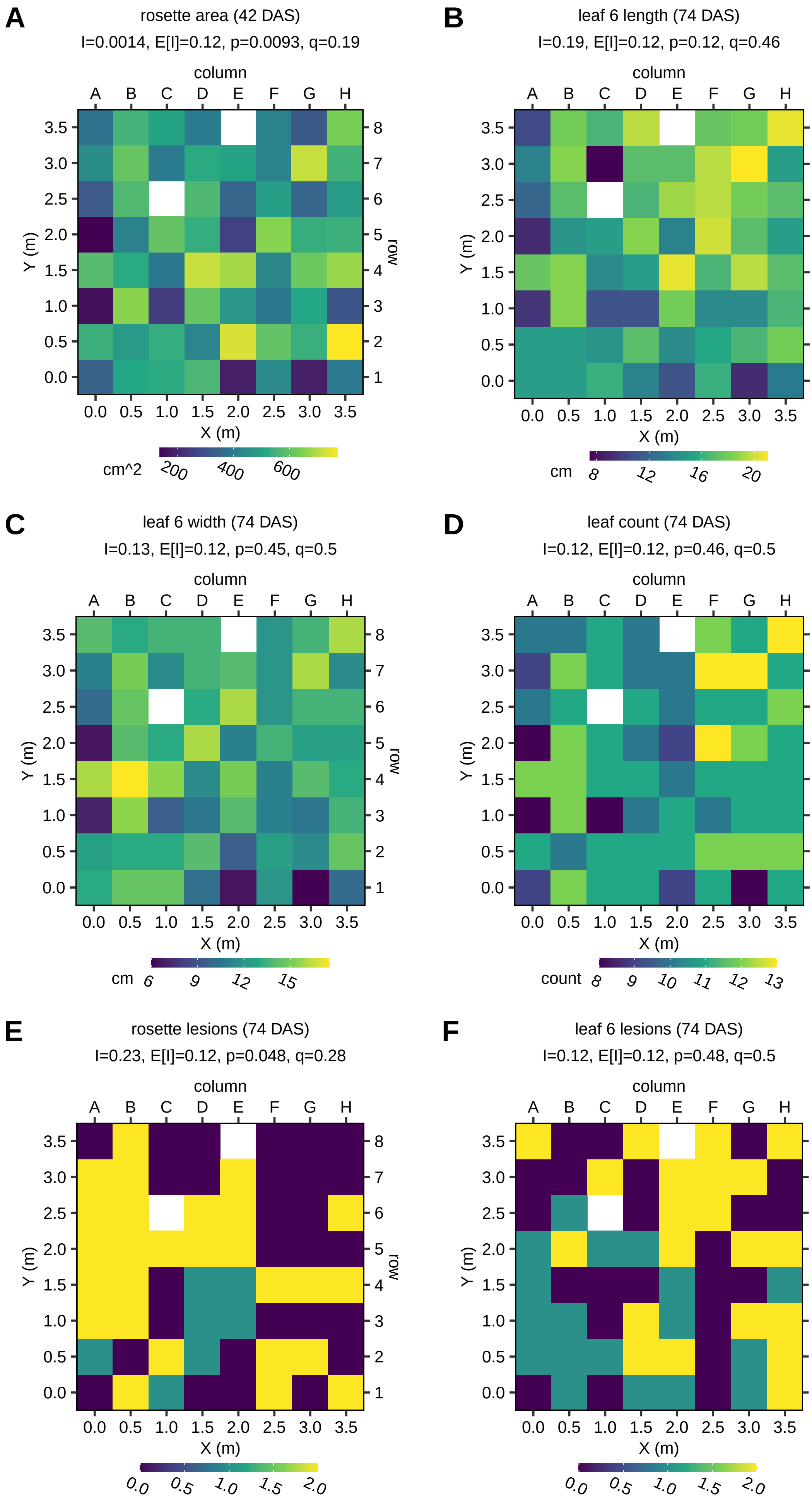
A
rosette area (42 DAS)
I=0.0014, E[I]=0.12, p=0.0093, q=0.19
cm^2 200 400 600
B
leaf 6 length (74 DAS)
I=0.19, E[I]=0.12, p=0.12, q=0.46
cm 8 12 16 20
C
leaf 6 width (74 DAS)
I=0.13, E[I]=0.12, p=0.45, q=0.5
cm 6 9 12 15
D
leaf count (74 DAS)
I=0.12, E[I]=0.12, p=0.46, q=0.5
count 8 9 10 11 12 13
E
rosette lesions (74 DAS)
I=0.23, E[I]=0.12, p=0.048, q=0.28
0.0 0.5 1.0 1.5 2.0
F
leaf 6 lesions (74 DAS)
I=0.12, E[I]=0.12, p=0.48, q=0.5
0.0 0.5 1.0 1.5 2.0
column
A B C D E F G H
row
1 2 3 4 5 6 7 8
X (m)
Y (m)
0.0 0.5 1.0 1.5 2.0 2.5 3.0 3.5

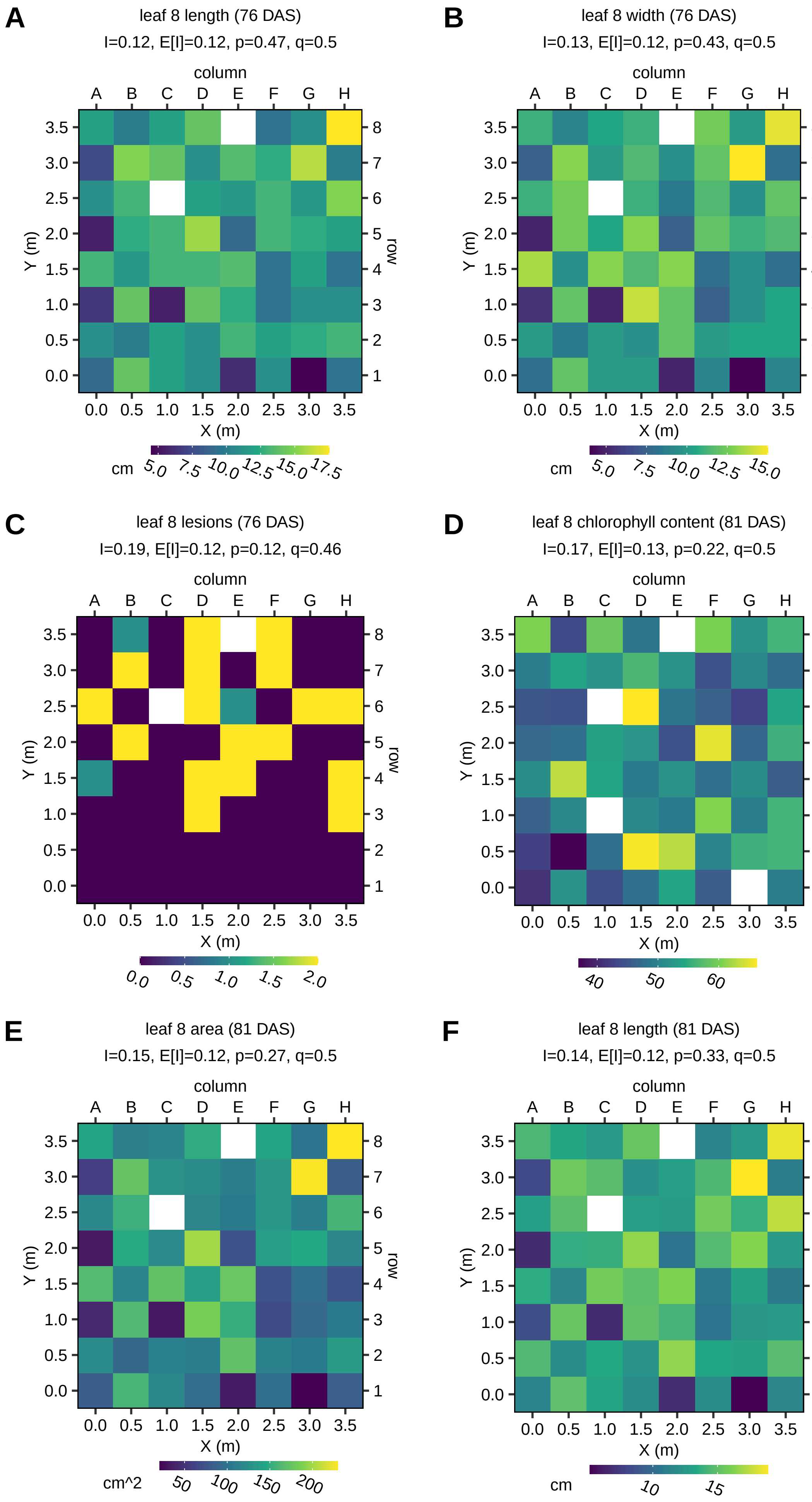
A
leaf 8 length (76 DAS)
I=0.12, E[I]=0.12, p=0.47, q=0.5
B
leaf 8 width (76 DAS)
I=0.13, E[I]=0.12, p=0.43, q=0.5
C
leaf 8 lesions (76 DAS)
I=0.19, E[I]=0.12, p=0.12, q=0.46
D
leaf 8 chlorophyll content (81 DAS)
I=0.17, E[I]=0.13, p=0.22, q=0.5
E
leaf 8 area (81 DAS)
I=0.15, E[I]=0.12, p=0.27, q=0.5
F
leaf 8 length (81 DAS)
I=0.14, E[I]=0.12, p=0.33, q=0.5
column
A B C D E F G H
Y (m)
X (m)
row
cm
cm^2

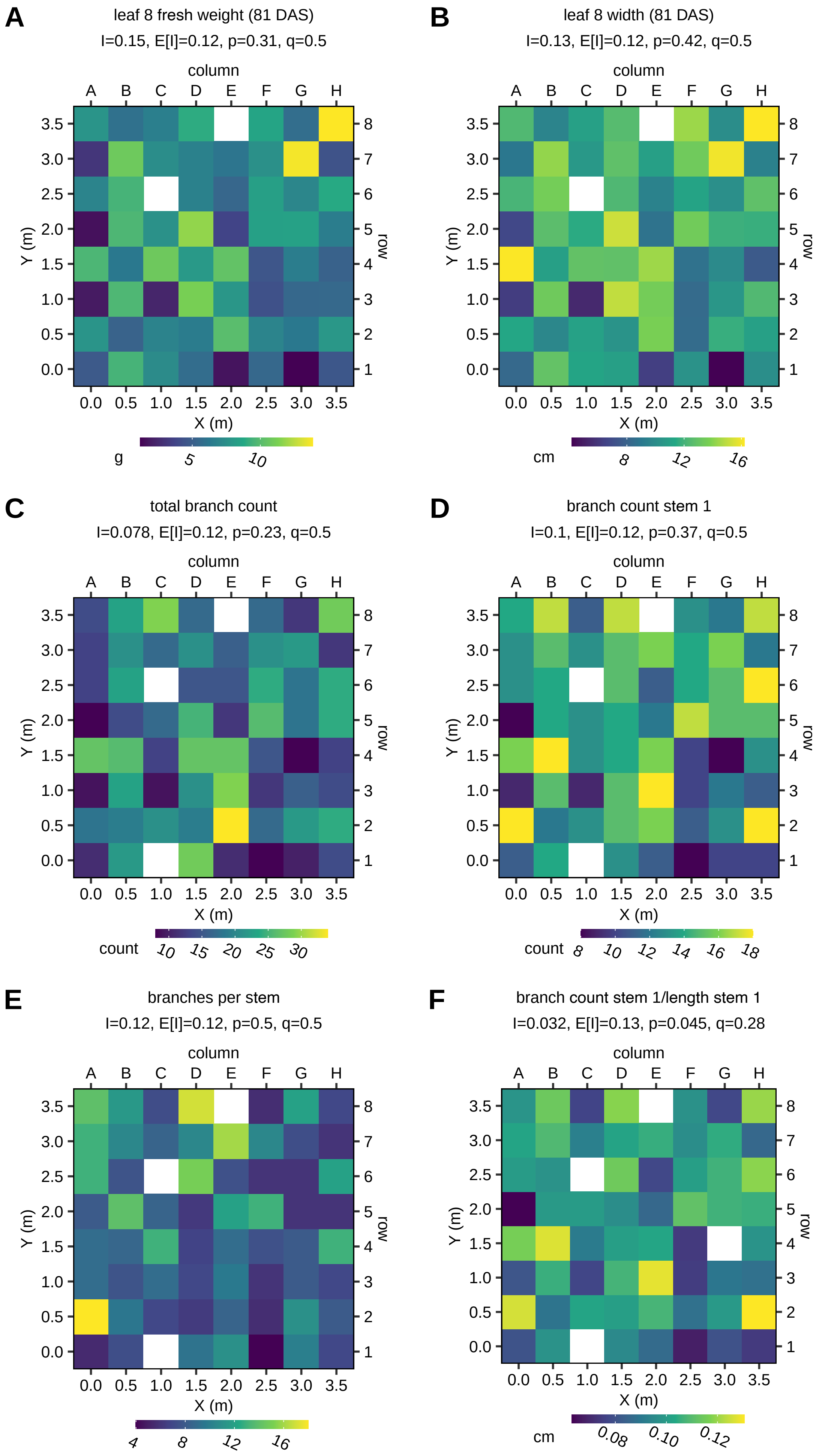
A
leaf 8 fresh weight (81 DAS)
I=0.15, E[I]=0.12, p=0.31, q=0.5
B
leaf 8 width (81 DAS)
I=0.13, E[I]=0.12, p=0.42, q=0.5
C
total branch count
I=0.078, E[I]=0.12, p=0.23, q=0.5
D
branch count stem 1
I=0.1, E[I]=0.12, p=0.37, q=0.5
E
branches per stem
I=0.12, E[I]=0.12, p=0.5, q=0.5
F
branch count stem 1/length stem 1
I=0.032, E[I]=0.13, p=0.045, q=0.28
column
row
X (m)
Y (m)
g
cm
count

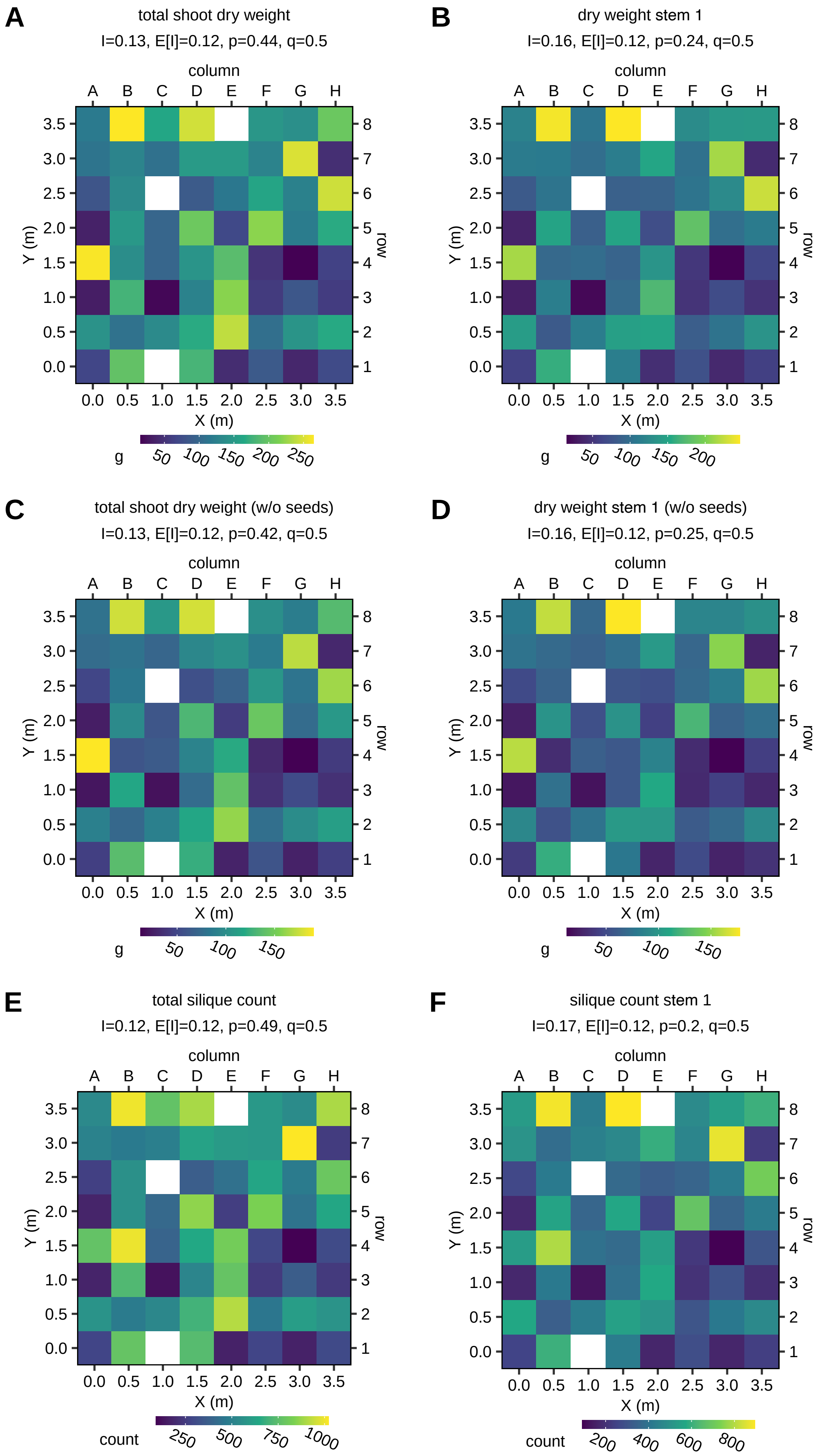
A
total shoot dry weight
I=0.13, E[I]=0.12, p=0.44, q=0.5
column
A B C D E F G H
3.5 3.0 2.5 2.0 1.5 1.0 0.5 0.0
Y (m)
8 7 6 5 4 3 2 1
row
0.0 0.5 1.0 1.5 2.0 2.5 3.0 3.5
X (m)
g 50 100 150 200 250
B
dry weight stem 1
I=0.16, E[I]=0.12, p=0.24, q=0.5
g 50 100 150 200
C
total shoot dry weight (w/o seeds)
I=0.13, E[I]=0.12, p=0.42, q=0.5
g 50 100 150
D
dry weight stem 1 (w/o seeds)
I=0.16, E[I]=0.12, p=0.25, q=0.5
g 50 100 150
E
total silique count
I=0.12, E[I]=0.12, p=0.49, q=0.5
count 250 500 750 1000
F
silique count stem 1
I=0.17, E[I]=0.12, p=0.2, q=0.5
count 200 400 600 800

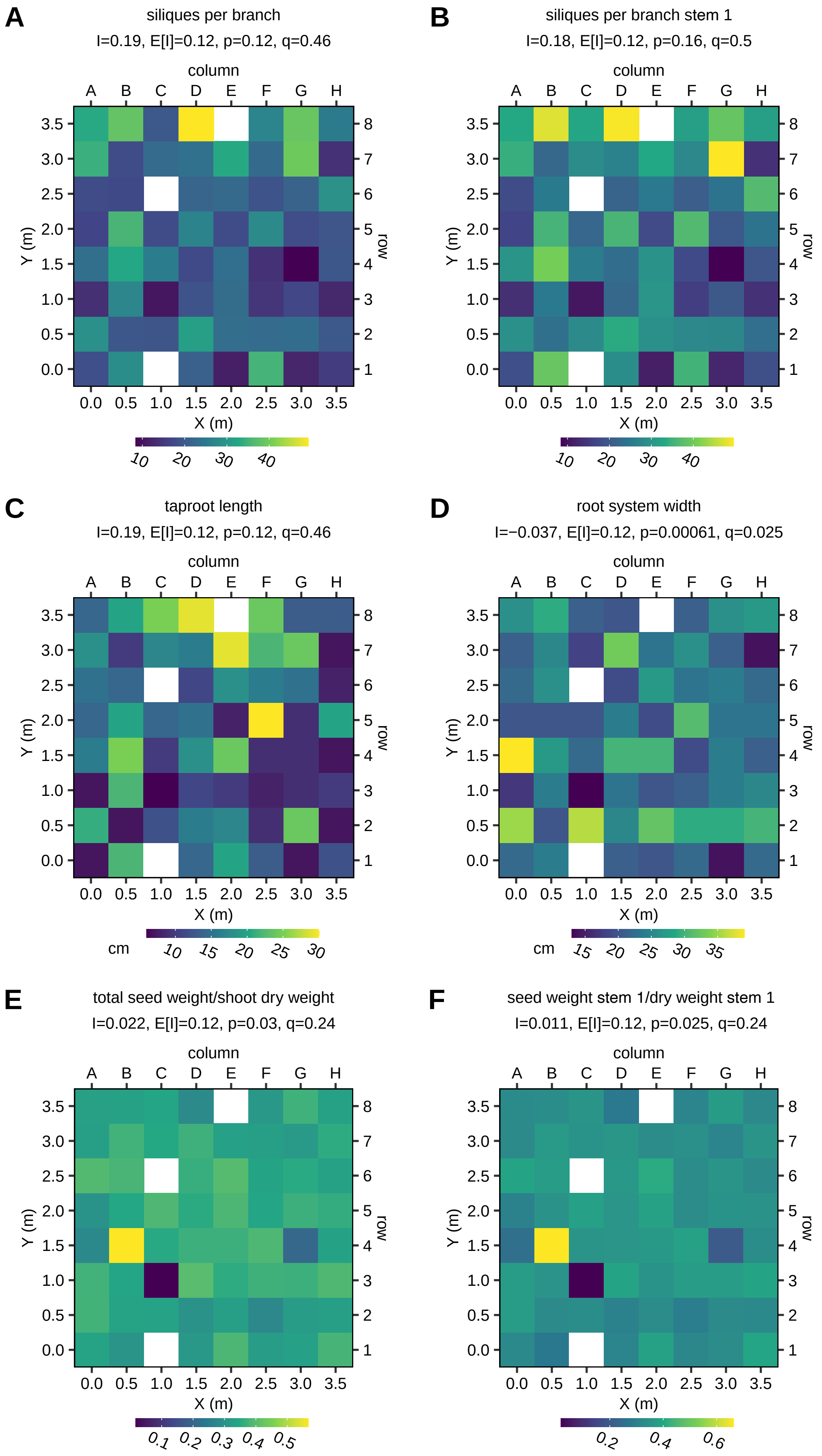
A
siliques per branch
I=0.19, E[I]=0.12, p=0.12, q=0.46
B
siliques per branch stem 1
I=0.18, E[I]=0.12, p=0.16, q=0.5
C
taproot length
I=0.19, E[I]=0.12, p=0.12, q=0.46
cm
D
root system width
I=−0.037, E[I]=0.12, p=0.00061, q=0.025
cm
E
total seed weight/shoot dry weight
I=0.022, E[I]=0.12, p=0.03, q=0.24
F
seed weight stem 1/dry weight stem 1
I=0.011, E[I]=0.12, p=0.025, q=0.24
column
A B C D E F G H
Y (m)
X (m)
row

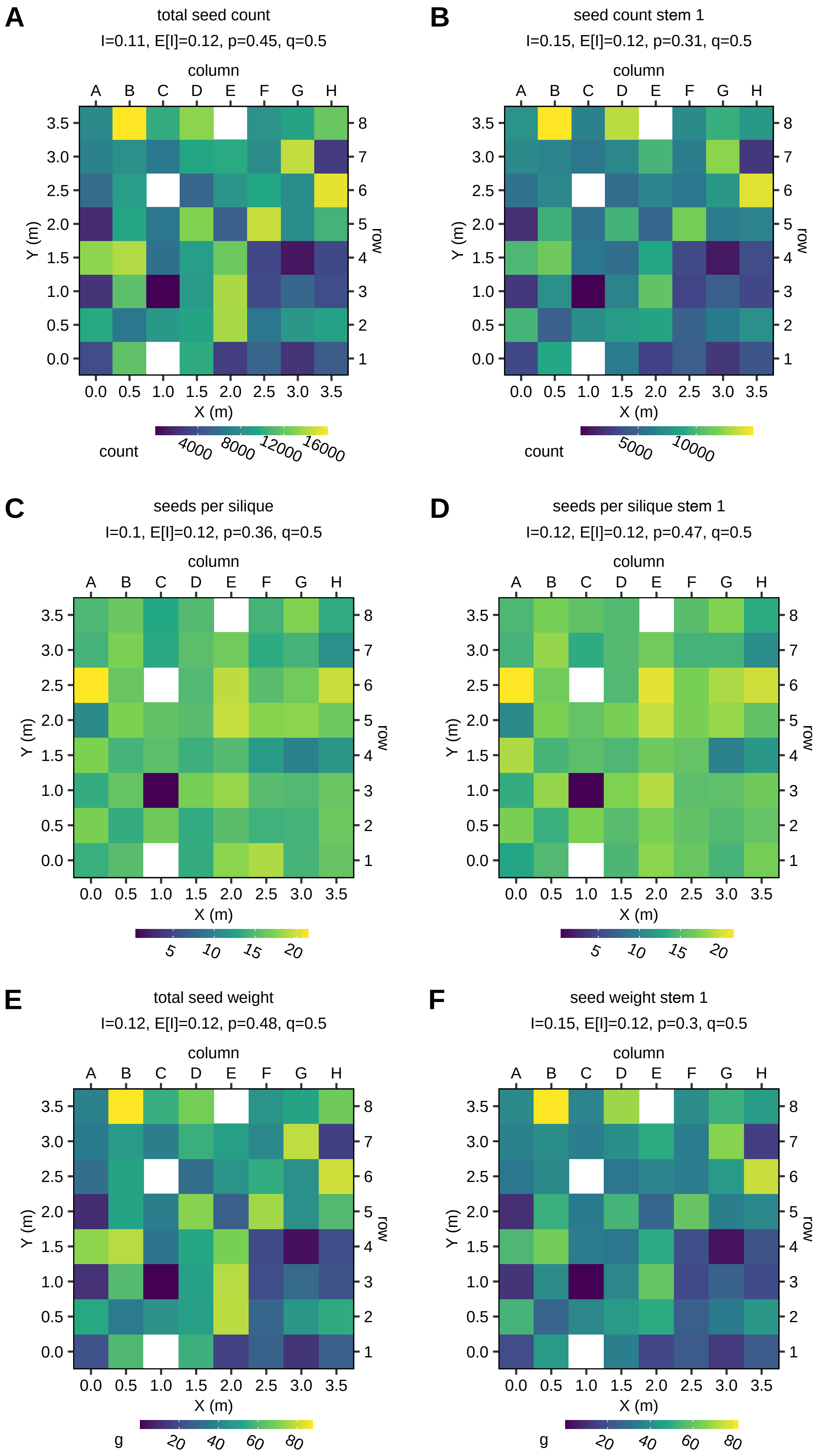
A
total seed count
I=0.11, E[I]=0.12, p=0.45, q=0.5
column
A B C D E F G H
3.5 3.0 2.5 2.0 1.5 1.0 0.5 0.0
Y (m)
8 7 6 5 4 3 2 1
row
0.0 0.5 1.0 1.5 2.0 2.5 3.0 3.5
X (m)
count
4000 8000 12000 16000
B
seed count stem 1
I=0.15, E[I]=0.12, p=0.31, q=0.5
column
A B C D E F G H
Y (m)
row
X (m)
count
5000 10000
C
seeds per silique
I=0.1, E[I]=0.12, p=0.36, q=0.5
column
A B C D E F G H
Y (m)
row
X (m)
5 10 15 20
D
seeds per silique stem 1
I=0.12, E[I]=0.12, p=0.47, q=0.5
column
A B C D E F G H
Y (m)
row
X (m)
5 10 15 20
E
total seed weight
I=0.12, E[I]=0.12, p=0.48, q=0.5
column
A B C D E F G H
Y (m)
row
X (m)
g
20 40 60 80
F
seed weight stem 1
I=0.15, E[I]=0.12, p=0.3, q=0.5
column
A B C D E F G H
Y (m)
row
X (m)
g
20 40 60 80

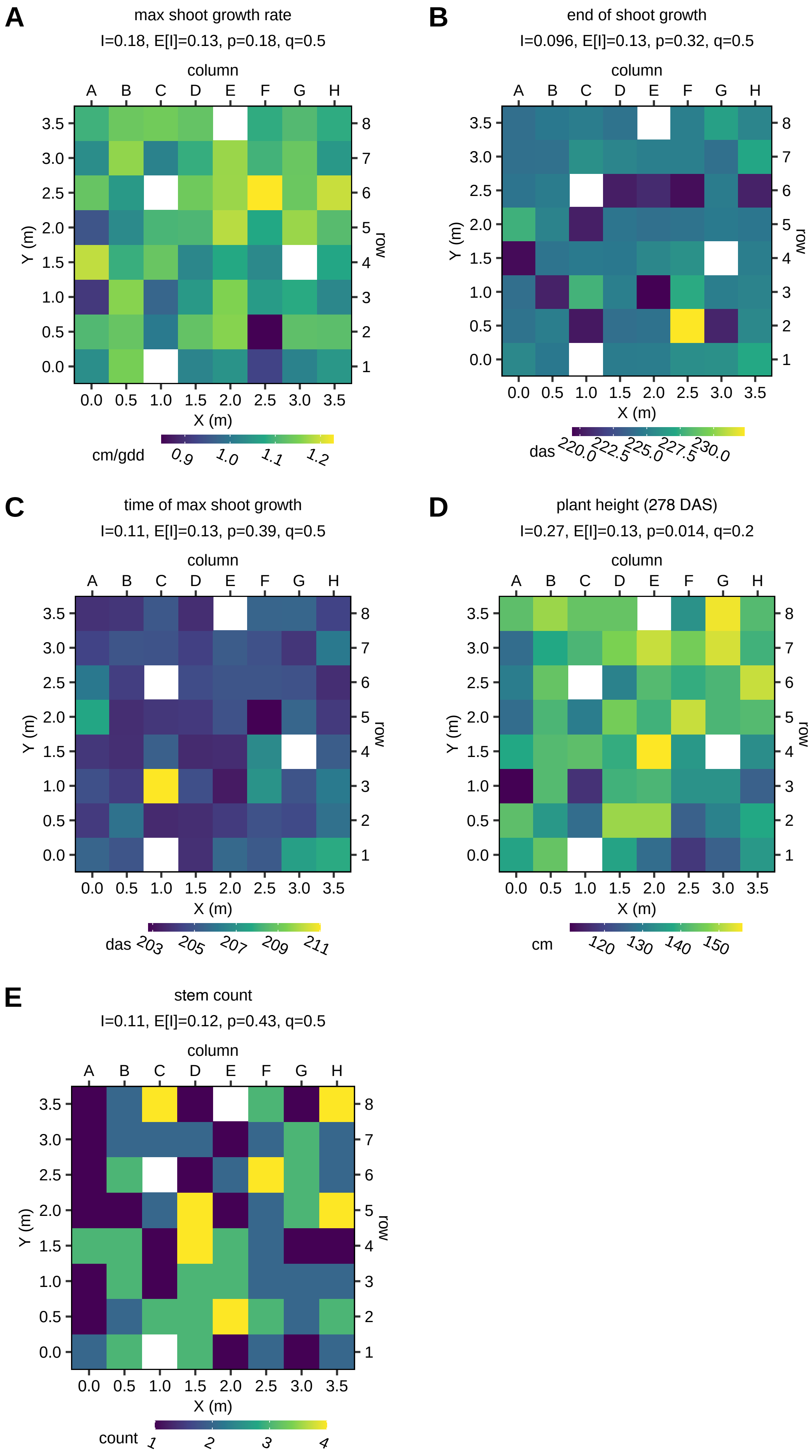
A
max shoot growth rate
I=0.18, E[I]=0.13, p=0.18, q=0.5
cm/gdd
B
end of shoot growth
I=0.096, E[I]=0.13, p=0.32, q=0.5
das
C
time of max shoot growth
I=0.11, E[I]=0.13, p=0.39, q=0.5
das
D
plant height (278 DAS)
I=0.27, E[I]=0.13, p=0.014, q=0.2
cm
E
stem count
I=0.11, E[I]=0.12, p=0.43, q=0.5
count
column
row
X (m)
Y (m)

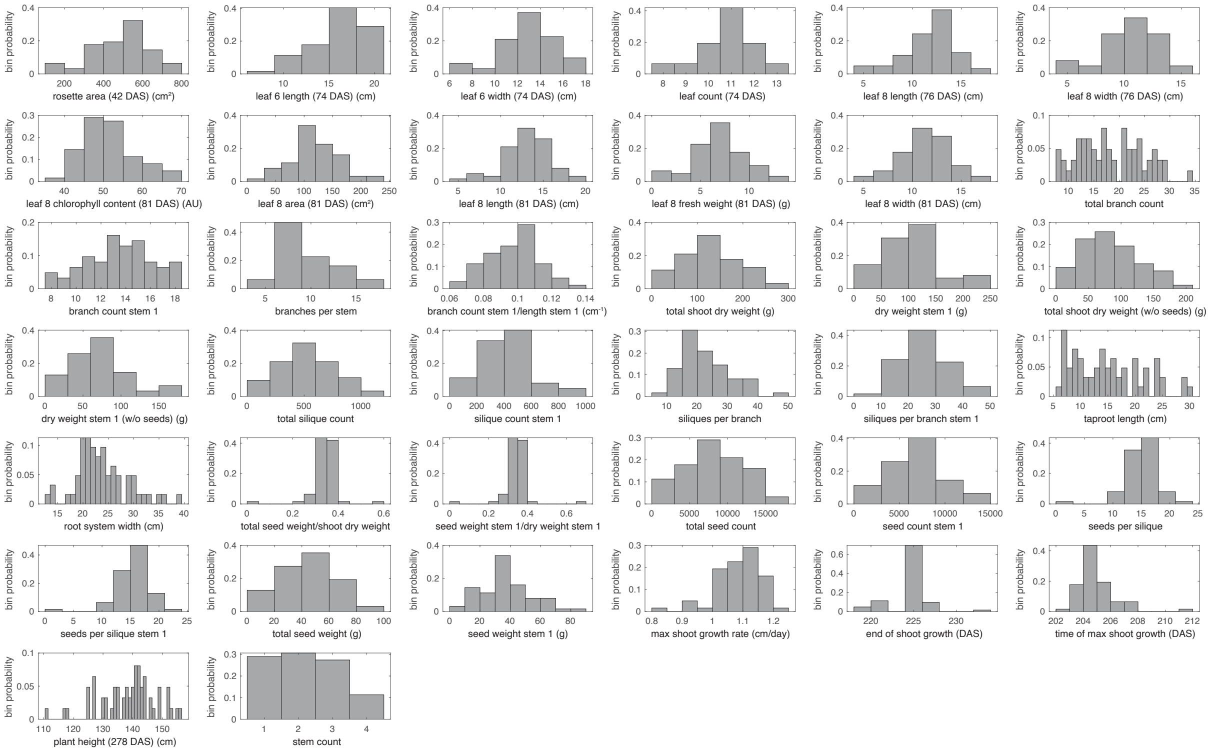

**Fig. S3** Phenotype histograms. Histograms were plotted using the 'histogram' function in Matlab R2018b, with automatic binning.

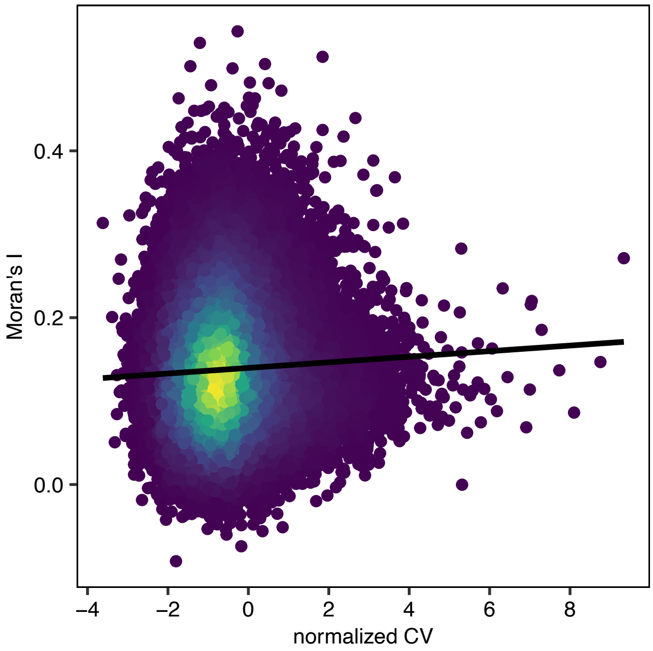


**Fig. S4.** Heatmap of Moran's I versus *normCV* values of gene expression profiles. Dots represent gene expression profiles and dot colors reflect dot density (yellow = high density, dark blue = low density). The line is an ordinary least-squares linear regression fit.

**Fig. S5** Phenotype predictions versus observations. Each plot shows the predicted versus measured values for the best-performing 'all genes' model for a given phenotype (Table 2). Qualitative and low-count phenotypes and phenotypes with median test $R^2$ values < 0 are not shown. Vertical grey lines range from the minimum to the maximum predicted value for a given plant across all model repeats, and colored dots represent predictions for the repeat with the median pooled $R^2$ score. Different marker colors indicate the 10 different test sets in this repeat. Perfect predictions are located on the dashed diagonal line in each panel.

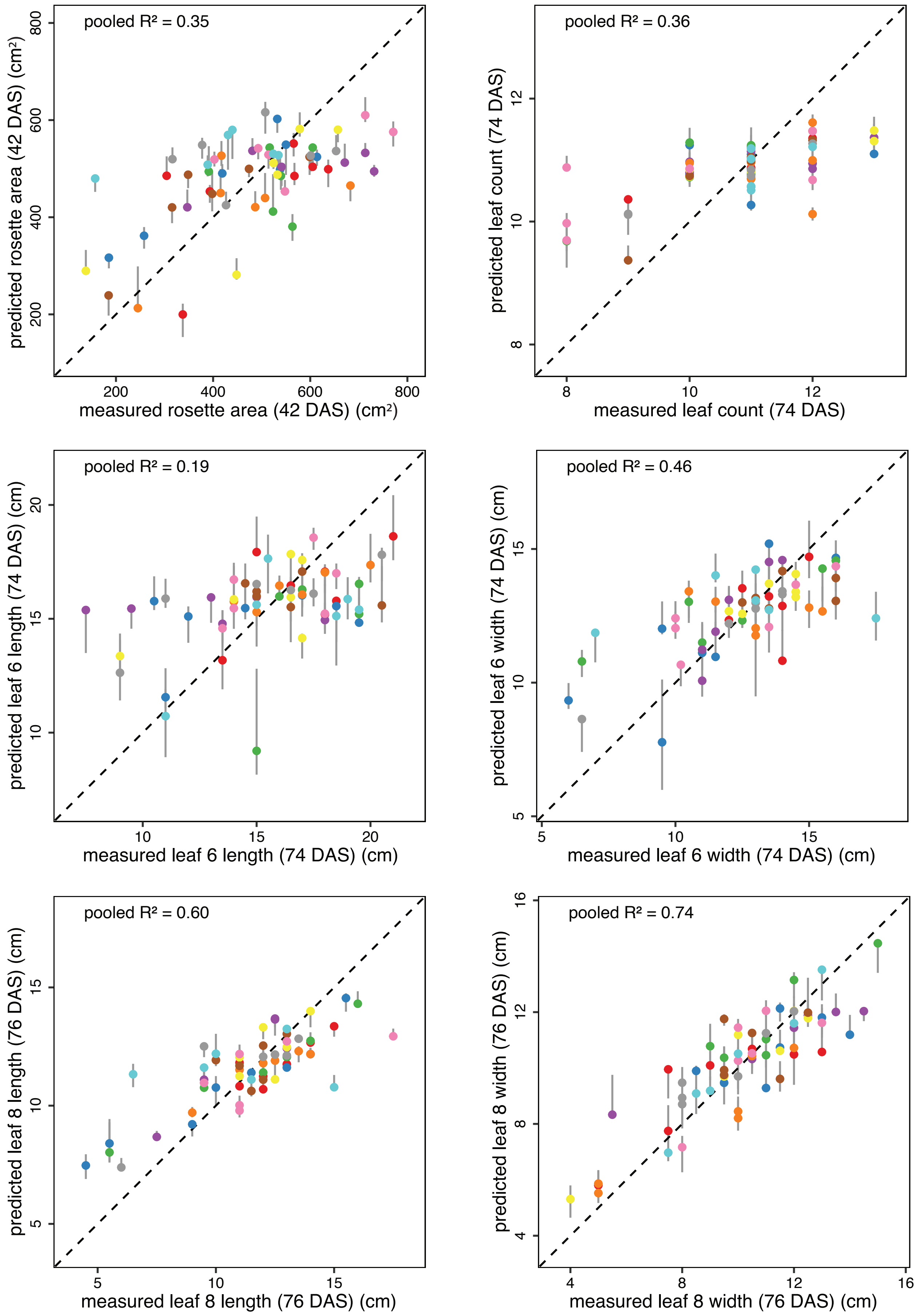

**Fig. S5 (continued).**

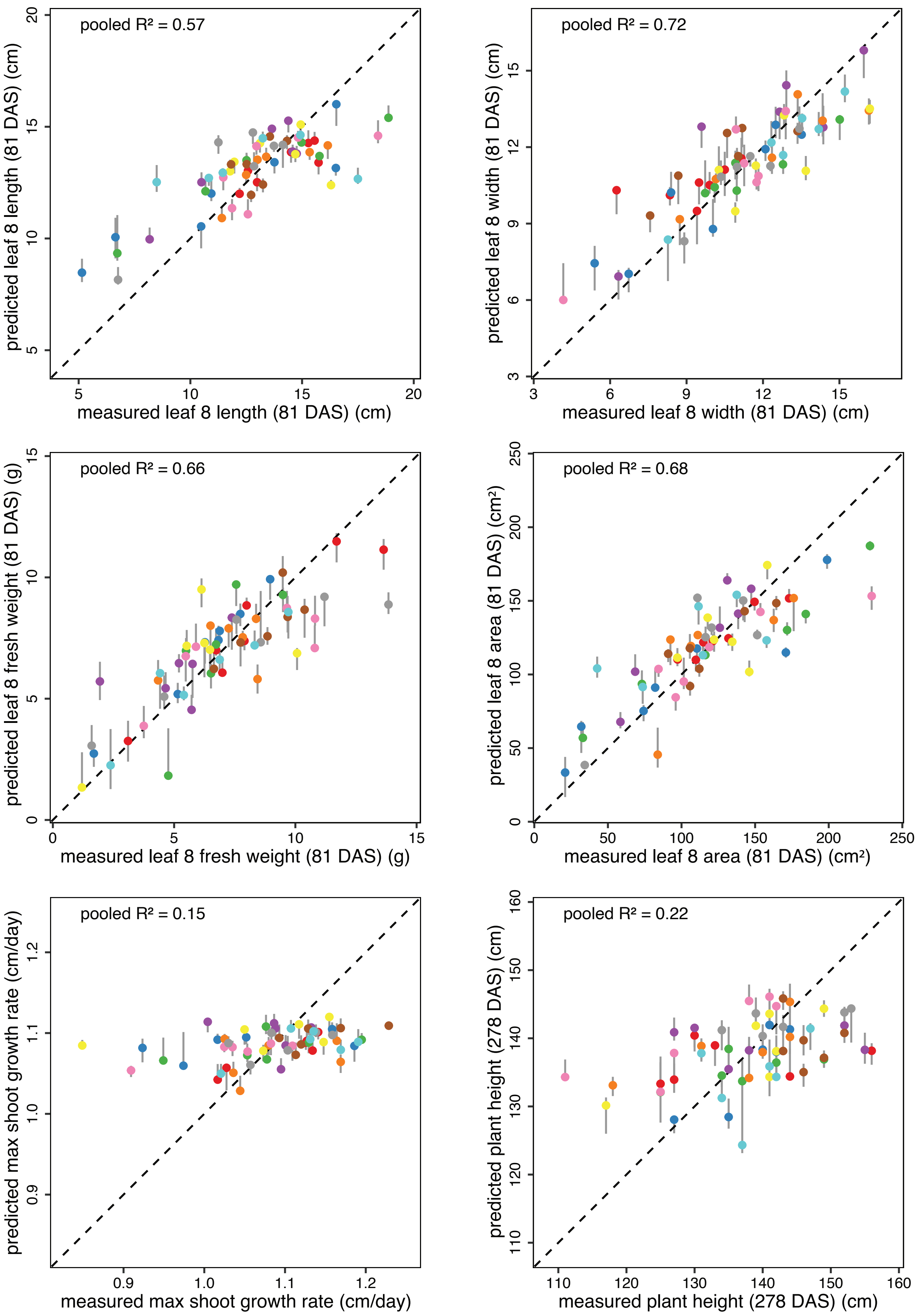

**Fig. S5 (continued).**

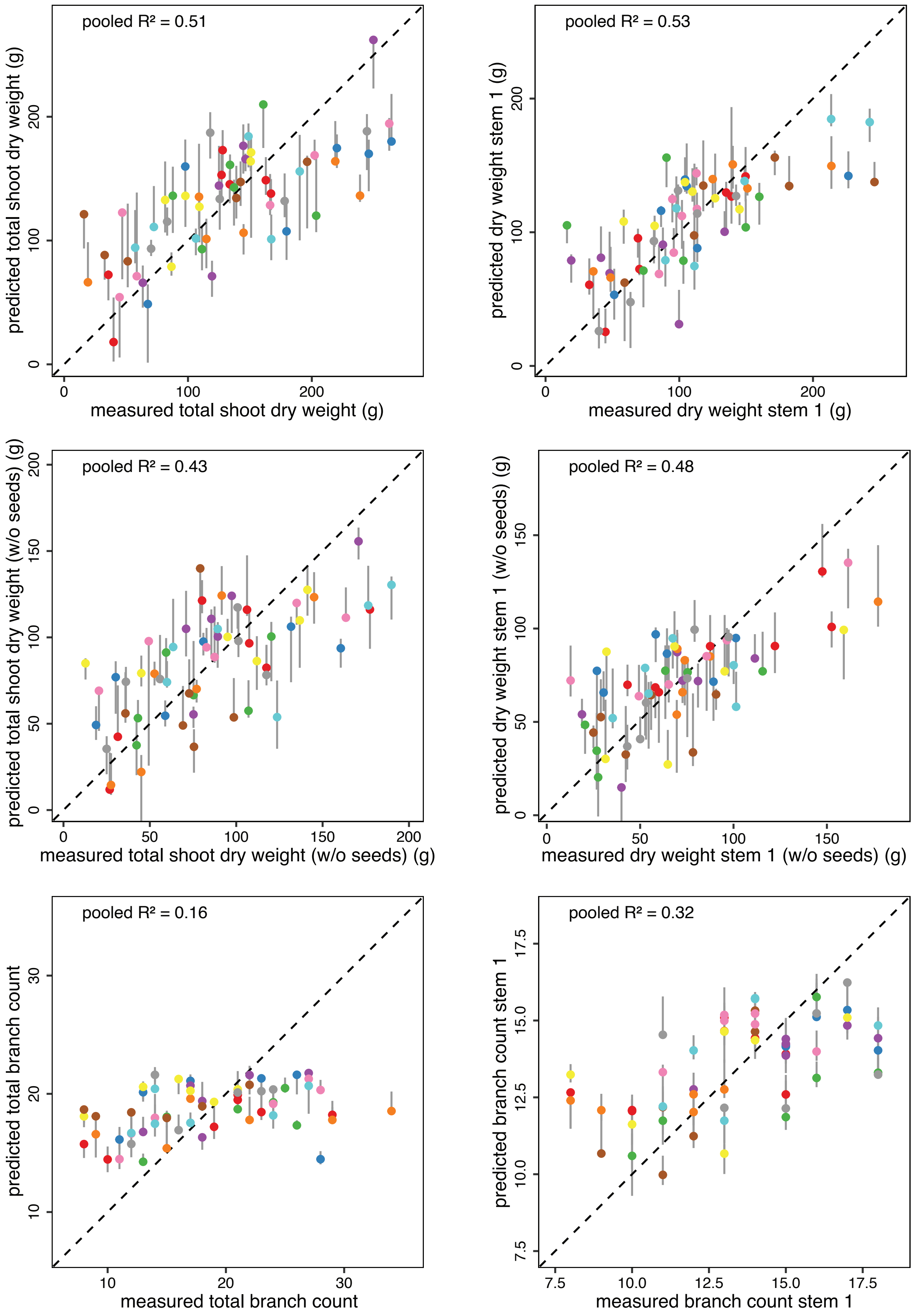

**Fig. S5 (continued).**

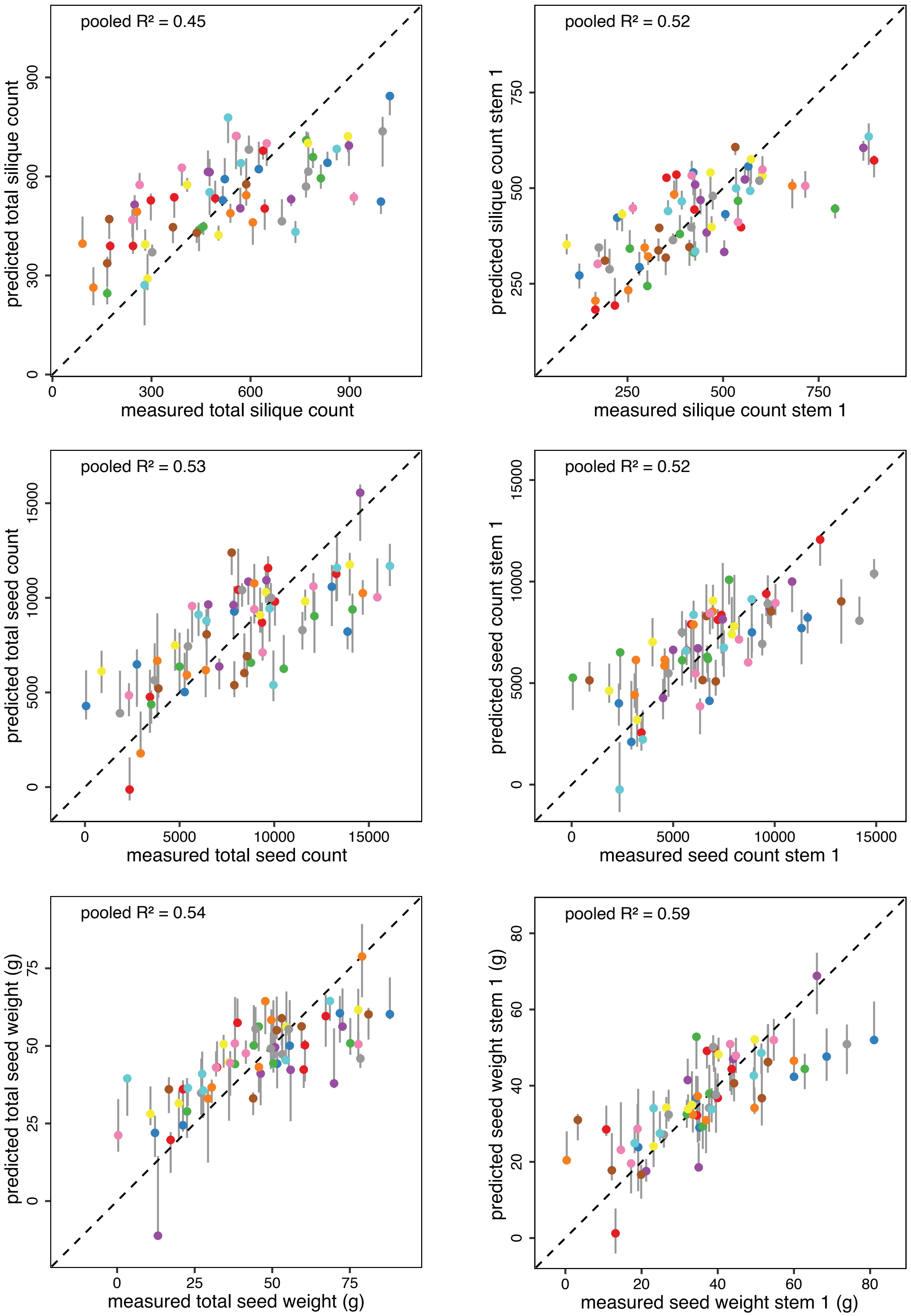

**Fig. S5 (continued).**

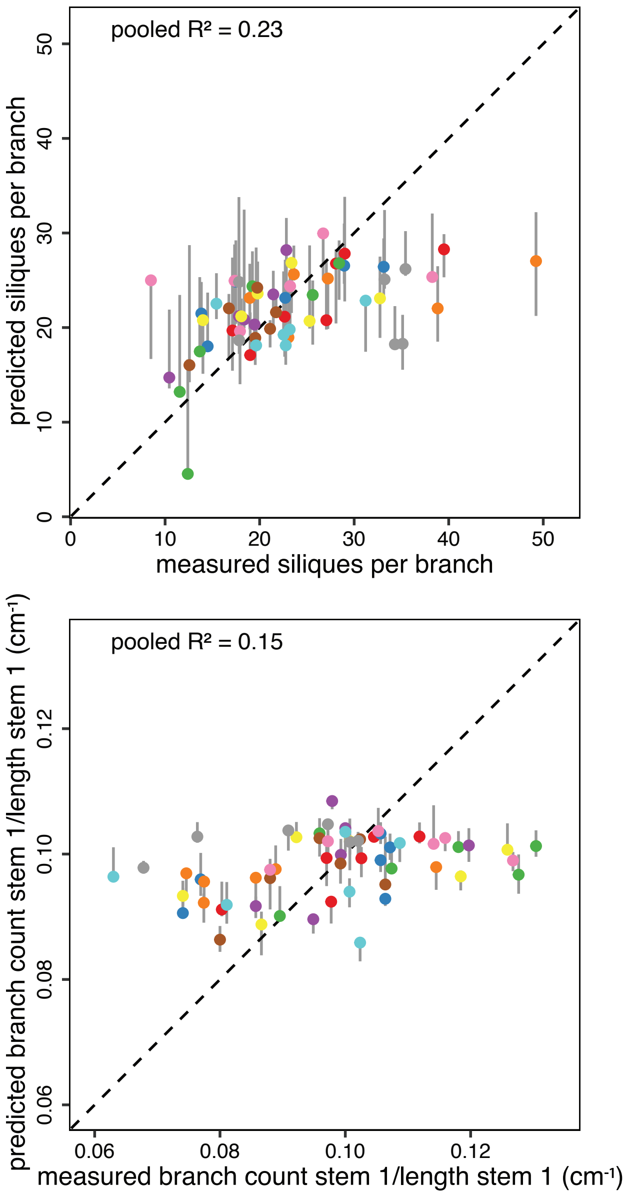

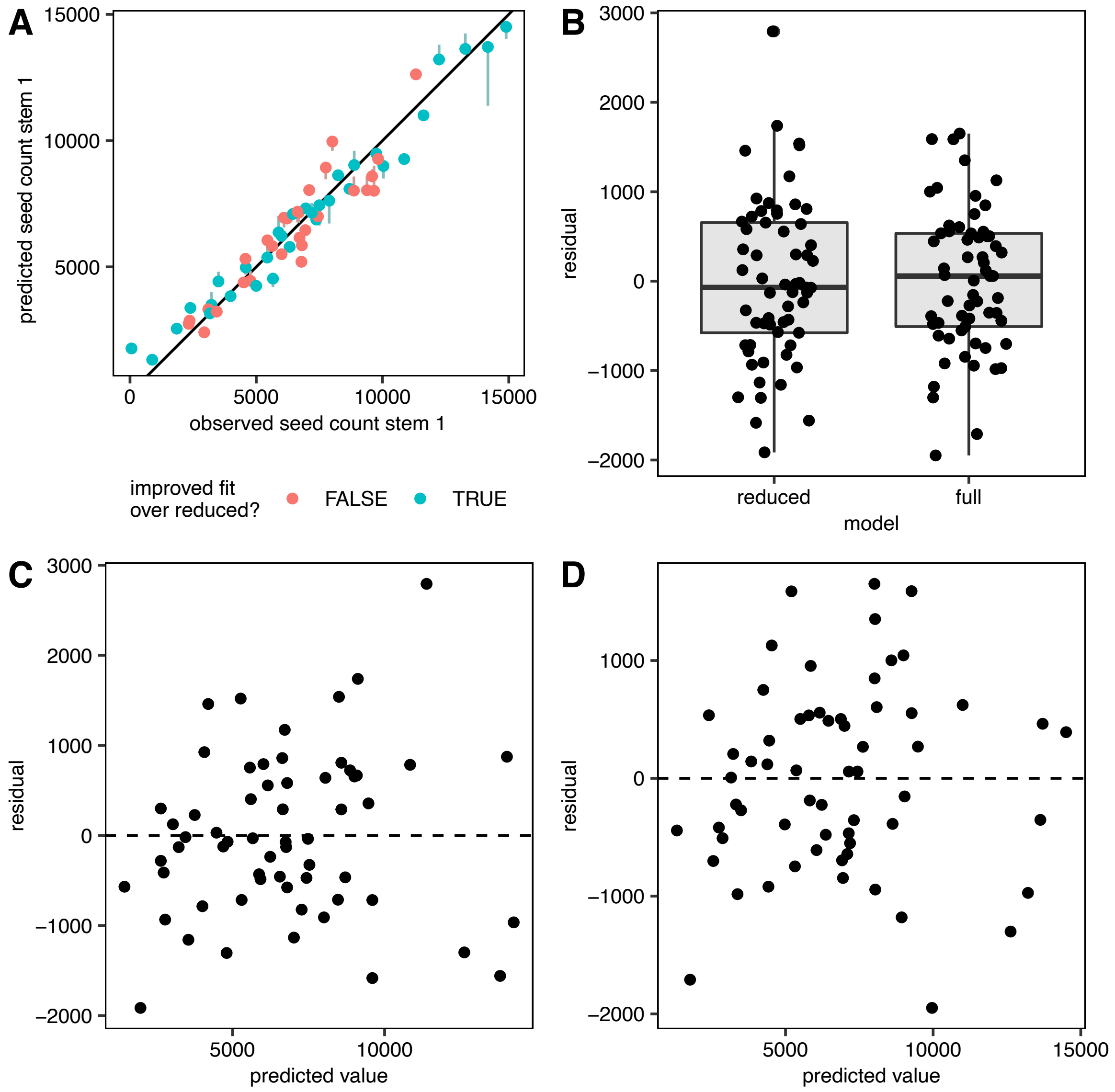


**Fig. S6** Performance of log-link model predicting seed count stem 1 conditioned on silique count stem 1 as a function of expression of the best predictor gene, *BnaCnng56980D*. **A** Plot of predicted versus observed seed counts on stem 1. Values predicted by the full model with heteroscedastic error structure are shown as dots. The ends of the tails attached to the dots indicate the phenotype values predicted by the reduced model (without gene expression effect). Blue dots indicate improved predictions in the full model versus the reduced model, red dots indicate worse predictions in the full model. **B** Distribution of residuals in the full and reduced models. **C** Residuals versus predicted values for the reduced model. **D** Residuals versus predicted values for the full model.

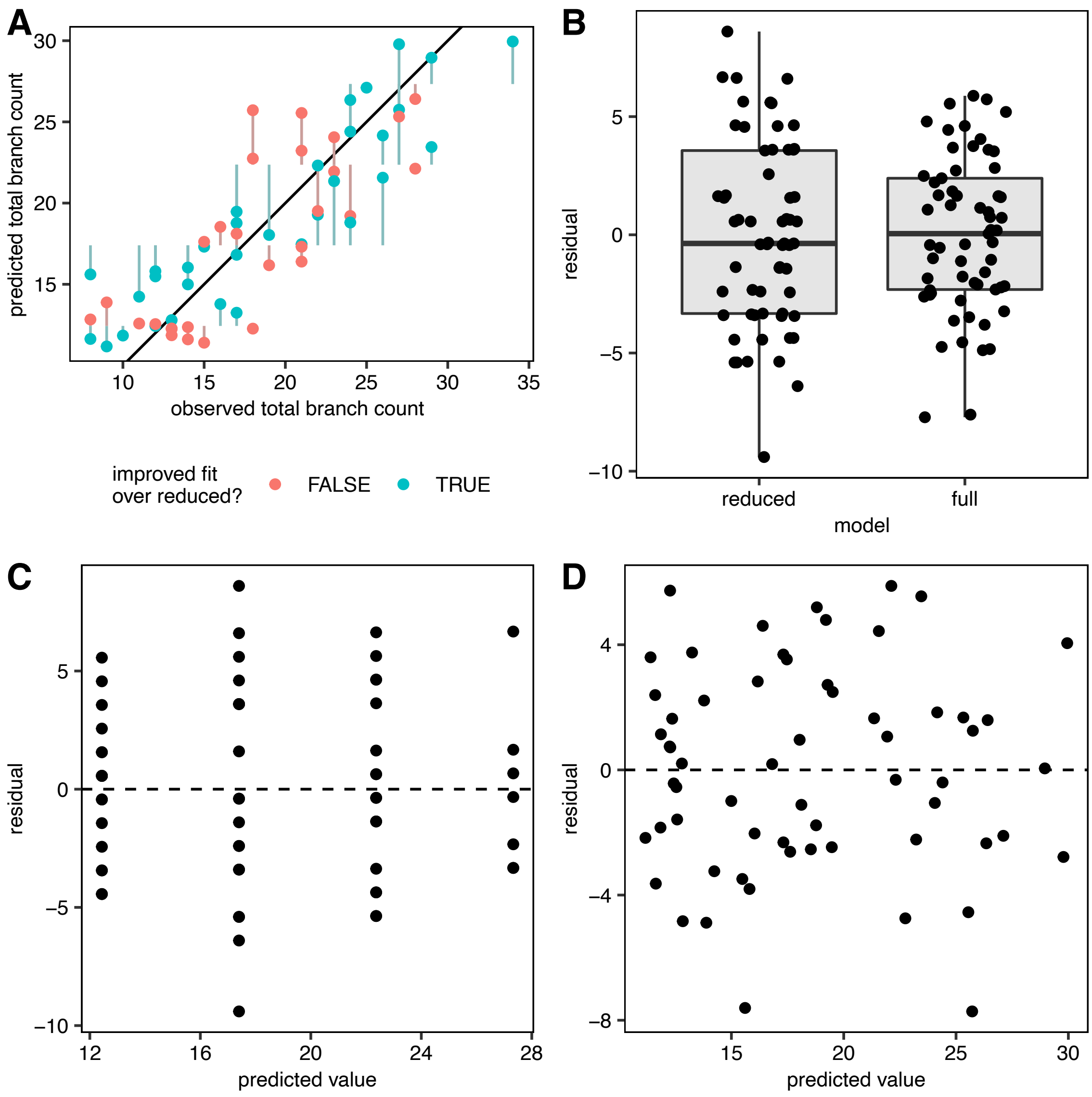


**Fig. S7** Performance of log-link model predicting total branch count conditioned on stem count as a function of expression of the best predictor gene, *BnaC01g26820D*. **A** Plot of predicted versus observed total branch counts. Values predicted by the full model with constant error variance are shown as dots. The ends of the tails attached to the dots indicate the phenotype values predicted by the reduced model (without gene expression effect). Blue dots indicate improved predictions in the full model versus the reduced model, red dots indicate worse predictions in the full model. **B** Distribution of residuals in the full and reduced models. **C** Residuals versus predicted values for the reduced model. Note that the predictions can only take a limited number of discrete values as predictions only depend on the stem count in the reduced model. **D** Residuals versus predicted values for the full model.

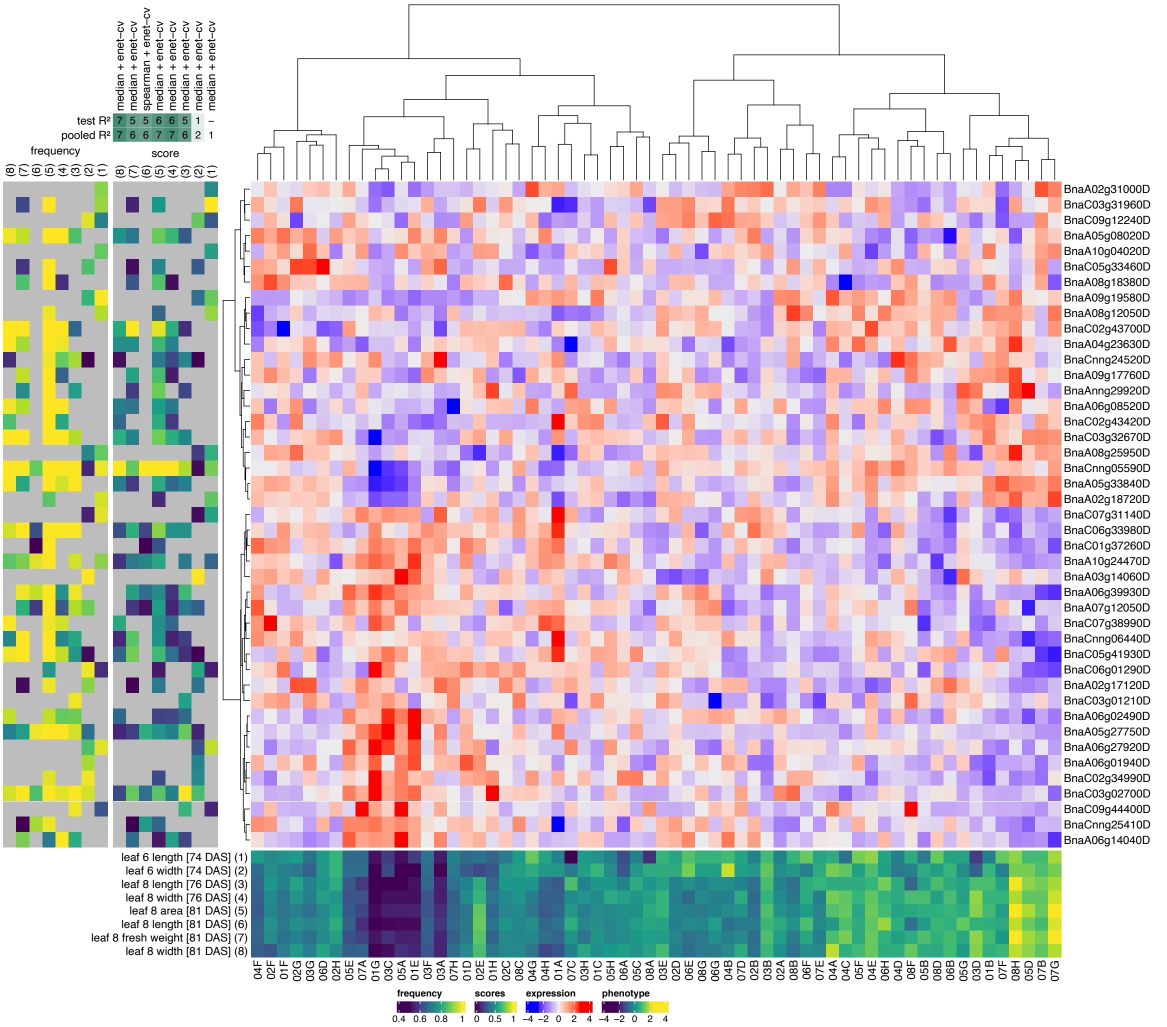


**Fig. S8** Top predictor genes in elastic net models of leaf phenotypes. A clustered heatmap of the z-scored gene expression profiles of the top genes for predicting leaf phenotypes is shown centrally (blue-red color scale, Ward.D2 hierarchical clustering). The leaf phenotypes concerned and their z-scored profiles across plants are shown at the bottom (dark blue-yellow heatmap with plant identifiers at the bottom). For each of these phenotypes, the top-10 most important genes (highest median elastic net coefficients across all 90 cross-validation splits) of the enet model with the highest median test $R^2$ score are included on the figure (gene identifiers are shown at right). The green-blue score panel to the left of the expression heatmap shows the median elastic net coefficients of the selected genes in each of the selected phenotype models, normalized to the maximum coefficient per model to make the color scales of the different models (columns) comparable. The mostly yellow frequency panel to the left of the score panel shows the frequencies at which genes were selected as features across all 90 cross-validation splits of a given model. Grey squares in the score and frequency panels indicate that a given gene was not selected as a feature in a given model. The phenotypes in the score and frequency panels are identified by numbers (1-8) on top of the panels, corresponding to the numbers associated with the phenotypes in the bottom phenotype panel. On top of the score panel, the feature selection techniques used in the best-scoring enet models for each phenotype are shown (median = selection of features with median *rlog* gene expression > 0, spearman = Spearman correlation, hsic-5000 = HSIC lasso, see Methods), as well as the corresponding test and pooled $R^2$ scores rounded to the nearest 0.1 and then multiplied by ten (e.g. a test $R^2$ score of 0.38 would be denoted as 4).

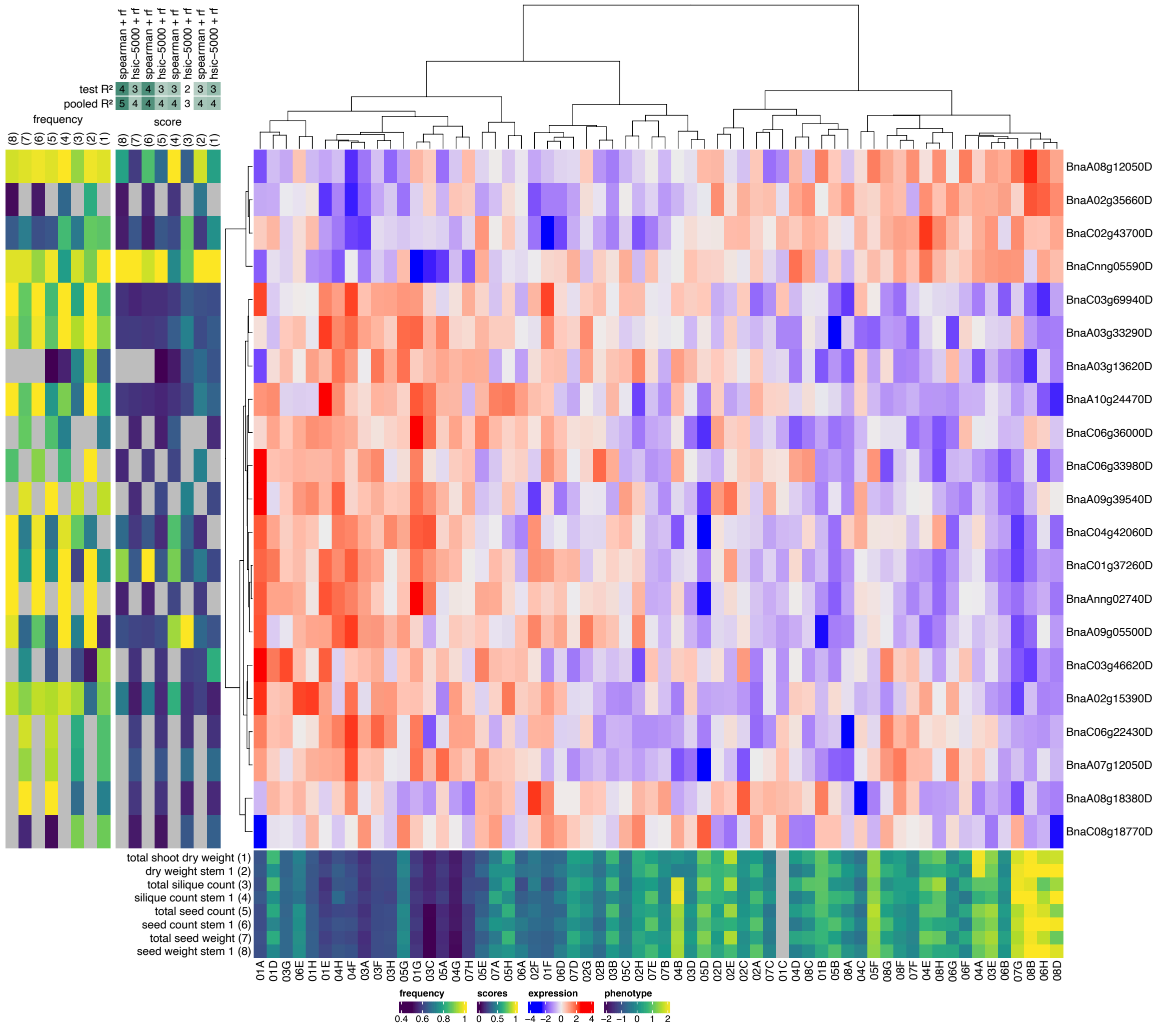


**Fig. S9** Top predictor genes in random forest models of yield phenotypes. A clustered heatmap of the z-scored gene expression profiles of the top genes for predicting yield phenotypes is shown centrally (blue-red color scale, Ward.D2 hierarchical clustering). The yield phenotypes concerned and their z-scored profiles across plants are shown at the bottom (dark blue-yellow heatmap with plant identifiers at the bottom). For each of these phenotypes, the top-10 most important genes (highest median gini importance across all 90 cross-validation splits) of the RF model with the highest median test $R^2$ score are included on the figure (gene identifiers are shown at right). The mostly blue score panel to the left of the expression heatmap shows the median gini importance scores of the selected genes in each of the selected phenotype models, normalized to the maximum importance score per model to make the color scales of the different models (columns) comparable. The yellow-green frequency panel to the left of the score panel shows the frequencies at which genes were selected as features across all 90 cross-validation splits of a given model. Grey squares in the score and frequency panels indicate that a given gene was not selected as a feature in a given model. The phenotypes in the score and frequency panels are identified by numbers (1-8) on top of the panels, corresponding to the numbers associated with the phenotypes in the bottom phenotype panel. On top of the score panel, the feature selection techniques used in the best-scoring RF models for each phenotype are shown (median = selection of features with median *rlog* gene expression > 0, spearman = Spearman correlation, hsic-5000 = HSIC lasso, see Methods), as well as the corresponding test and pooled $R^2$ scores rounded to the nearest 0.1 and then multiplied by ten (e.g. a test $R^2$ score of 0.38 would be denoted as 4).

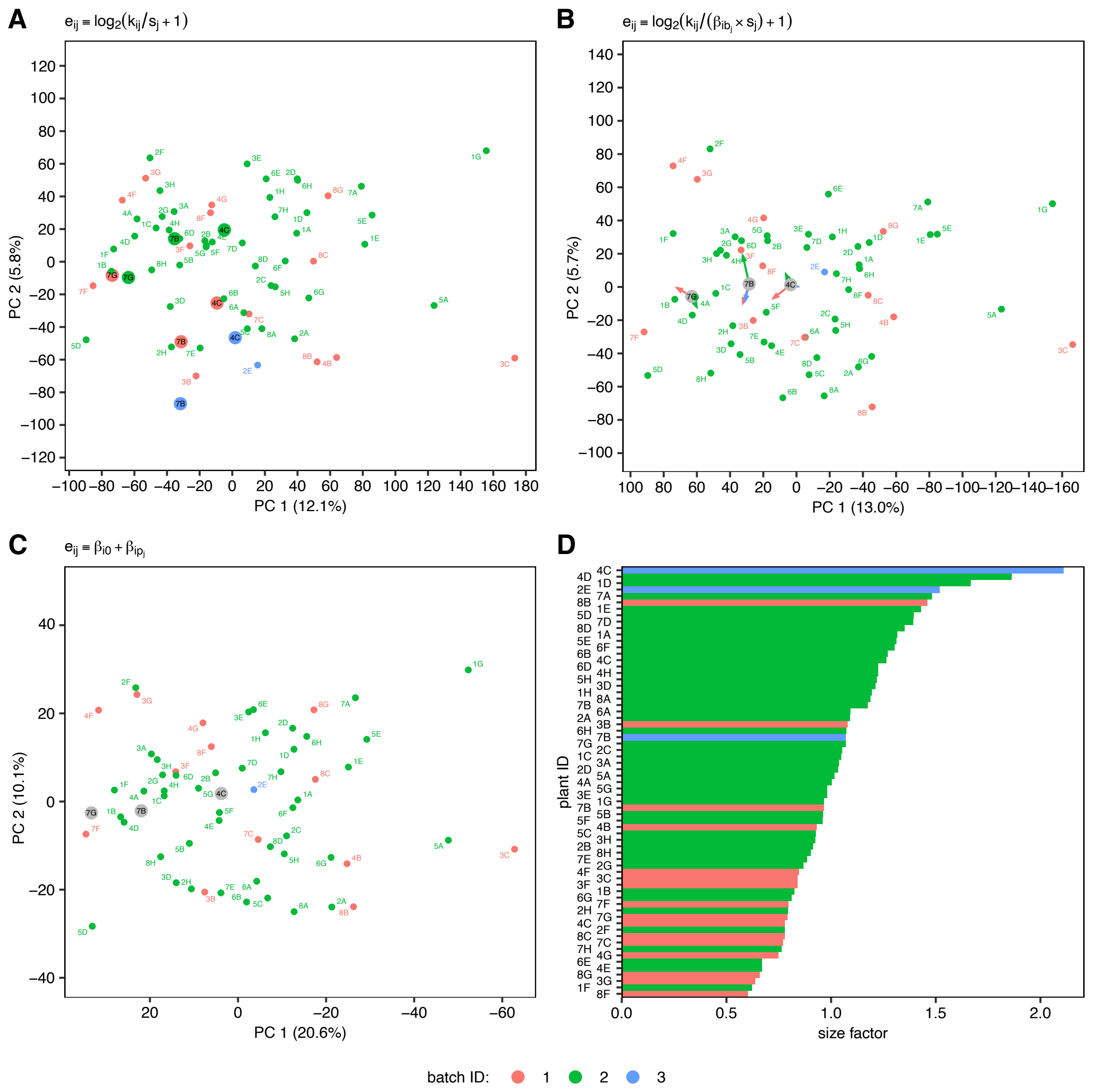


**Fig. S10.** Sequencing batch effects on RNA-seq count data. **A-C** Effect of RNA-seq batch on gene expression in principal component (PC) space. Samples are colored by batch ID and labeled with plant IDs. Samples included in several batches are indicated with bigger dots and black labels. **A** First two PCs of $\log_2$-tranformed library size-corrected expression profiles, before batch correction. Repeats of samples 4C and 7B are spaced far apart in PC 2, but align well on PC 1. **B** First two PCs of $\log_2$-transformed library size-corrected and batch-corrected expression profiles. Grey dots indicate corrected expression profiles that were averaged across repeats before log-transforming, and colored arrows point to the batch-specific positions of the samples concerned before averaging. Comparison with **A** shows that batch correction diminishes but does not completely eliminate the gene expression differences between sample repeats. The non-log-transformed version of the data in **B** was used for variance analysis (see Methods). **C** First two PCs of gene expression profiles obtained through the modified *rlog* transformation, which accounts for library size and batch effects and unites technical repeats in one estimate (grey dots, see Methods). **D** Size factors estimated by DESeq2 for each sample and batch.

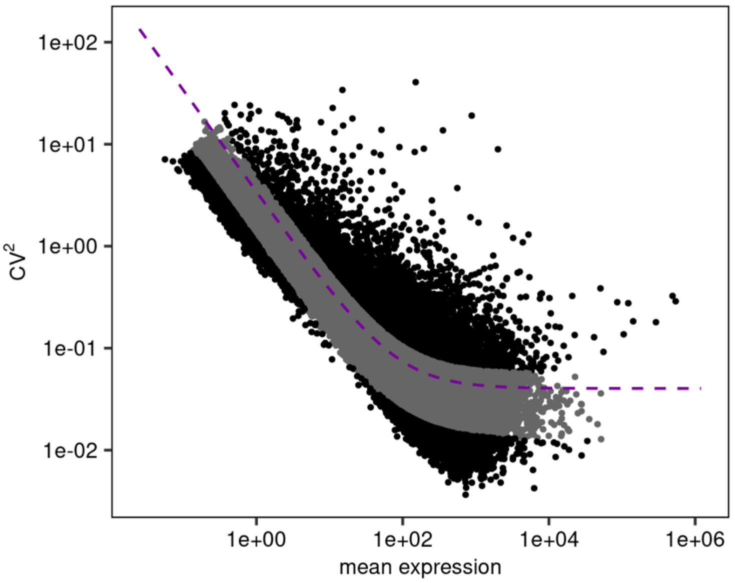


**Fig. S11. Gene expression variability in the *B. napus* single-plant dataset.** The squared CV is plotted versus the mean expression for genes expressed in ≥ 10 samples (dots). A fitted trendline (see Methods) is shown in purple, the top and bottom 10% of genes ranked by normalized CV (*normCV*, see Methods) are shown in black.
